# Retrospective Evaluation of Cyclosporine in the Treatment of Chronic Hepatitis in Dogs: 2010-2017

**DOI:** 10.1101/453977

**Authors:** Ullal Tarini, Ambrosini Yoko, Rao Sangeeta, Webster Cynthia RL, Twedt David

## Abstract

**Background:** Idiopathic chronic hepatitis (CH) in dogs is a prevalent hepatic disorder. The etiology is poorly understood; however, there is evidence to support an immune-mediated pathogenesis. No literature exists investigating the efficacy of cyclosporine (CsA) therapy for CH in dogs.

**Objectives:** To retrospectively evaluate the efficacy and adverse side effects of CsA in the treatment of CH in dogs, and to identify factors that impact response to CsA therapy.

**Animals:** 48 client-owned dogs diagnosed with CH treated with CsA for at least 2 weeks.

**Methods:** Retrospective review of medical records between the years 2010-2017.

**Results:** Twenty-two breeds of dogs were included of which 54% were spayed females, 42% neutered males and 4% intact males. Median age was 8.5 years (range, 0.7-14 years). Complete remission (normalization of alanine aminotransferase [ALT]) in response to CsA was attained in 79% of dogs (38/48). Median dose of CsA at the time of remission was 7.9 mg/kg/day (range, 2.5-12.7mg/kg/day) and median time to remission was 2.5 months (range, 0.75-18 months). None of the factors evaluated including clinical score, presence of ascites, hypoalbuminemia, hyperbilirubinemia, prolonged coagulation times, dose, or duration of therapy influenced remission. Common side effects were gastrointestinal signs in 38% (18/48) and gingival hyperplasia in 25% (12/48) of dogs.

**Conclusion and Clinical Importance:** CsA proved to be a tolerated and effective medication in attaining remission in dogs with idiopathic CH based on normalization of serum ALT. None of the evaluated factors were shown to negatively impact response to CsA or remission of disease.

---

Chronic hepatitis (CH) in dogs is a common and important hepatic disorder in dogs.^1–6^ The exact prevalence of the disease is unknown, but in one study, 0.5% of the referral population to a university hospital was diagnosed with primary hepatitis, of which 64% were diagnosed with CH.^6^ CH is defined by persistent ALT (alanine aminotransferase) elevations > 2 times normal in the presence of histopathological changes that include a mixed, primarily monocellular inflammatory cell infiltrate accompanied by evidence of hepatocyte cell death and variable amounts of fibrosis, ductular response and regenerative nodule formation.^7,8^ The specific etiology of the disease is poorly understood with a majority of cases labeled as idiopathic.^6^ A growing body of evidence, however, supports an immune-mediated pathogenesis in some dogs. This evidence includes abnormal MHC Class II protein expression on hepatocytes,^9–11^ presence of serum autoantibodies,^12–15^ lymphocytic infiltrates in the liver, ^6,16^ female and breed predispositions,^2,3,6,17–21^ association with other autoimmune diseases,^18^ and a positive response to immunosuppression.^20–22^ Two studies have suggested that there are dogs with CH that respond to immunosuppressive corticosteroids.^21,22^ However, corticosteroids in dogs can cause significant, often intolerable side effects (polyuria/polydipsia, panting, muscle atrophy, thrombosis, and gastrointestinal ulceration). ^23^ Corticosteroids also induce morphologic vacuolar change in hepatocytes and are associated with the induction of serum liver enzyme elevations.^23^ The latter makes the laboratory monitoring of response to therapy particularly challenging.

Cyclosporine (CsA) is an immunomodulatory medication that inhibits the activation and proliferation of T-lymphocytes.^24^ CsA has proven effective in dogs and humans in the management of various immune mediated disorders including atopic dermatitis, ^25–28^ immune mediated thrombocytopenia, ^29^ inflammatory bowel disease,^30^ immune-mediated polyarthritis,^31^ and renal transplantation.^32^ CsA is well tolerated in most dogs. The most common reported adverse side effects associated with CsA in dogs are gastrointestinal signs^25–28^ and less commonly, opportunistic infections,^33–37^ gingival hyperplasia,^38^ papillomatosis, and hirsuitism.^25–28^ Although prednisone and azathioprine are first line immunosuppressive therapies for immune hepatitis in humans, CsA had been successfully used as a second line therapy in patients that are refractory or intolerant of this drug combination.^39,40^ To date, no study has evaluated the use of CsA in the treatment of suspected immune mediated CH in dogs.

The primary aim of this study was to retrospectively evaluate the safety and efficacy of CsA in the treatment of CH. A secondary aim was to identify if any factors previously shown to have prognostic significance; such as, the presence of ascites,^6,41^ clinical score,^18^ hyperbilirubinemia,^6,42^ hypoalbuminemia,^6,21^ or prolongations in prothrombin time (PT)/activated partial thromboplastin time (aPTT),^6,18,21^ influenced CsA treatment response and biochemical remission.

## Materials and methods

### Case selection

Electronic medical records from Colorado State University Veterinary Teaching Hospital and Foster Hospital for Small Animals at Cummings School of Veterinary Medicine at Tufts University between the years 2010-2017 were reviewed. Dogs were included if they underwent an ultrasound guided, laparoscopic, or surgical liver biopsy and were diagnosed with CH by a board-certified veterinary pathologist. Patient presentation, diagnostic test results, liver histopathology reports, and treatment response to cyclosporine were reviewed by one of two board certified internists (DCT, CLW) to ensure adherence to WSAVA (World Small Animal Veterinary Association) histologic criteria and limit inclusion to dogs with presumed idiopathic CH.^8^ Dogs with exposure to chronic hepatotoxic drugs/supplements or evidence of systemic infectious disease were excluded. Dogs with elevated hepatic copper levels (> 1500 ug/g dry weight) were included only if hepatic copper reduction therapy with penicillamine, zinc and/or a copper restricted diet had not normalized ALT prior to initiation of CsA. Dogs that received antibiotics or concurrent immunosuppressive medications were included as long as the antimicrobial or immunomodulatory therapy had failed to achieve remission (normalization of ALT) prior to initiation of CsA. It was not required that any of the copper reduction, antimicrobial, or immunosuppressive therapies be administered for a specific duration of time. However, dogs that received less than 2 weeks of CsA treatment were excluded due to inability to evaluate medication efficacy with the short duration of treatment.

### Data collection

Data regarding signalment, body weight, clinical signs at presentation, presence of ascites (determined by abdominal ultrasound), hepatic encephalopathy (i.e. increased serum ammonia level or presence of diffuse neurologic signs that were responsive to lactulose, metronidazole/neomycin, and/or low protein diet), concurrent medications, diet, dose of CsA, possible side effects of CsA, prothrombin time (PT), activated partial thromboplastin time (aPTT), serum alkaline phosphatase (ALP), serum alanine aminotransferase, (ALT), serum aspartate aminotransferase (AST), serum gamma-glutamyl transferase, (GGT), serum albumin, serum total bilirubin, and serum globulin values at the time of initial presentation to the hospital were extracted from electronic medical records for each dog. Individual clinical scores (a numeric scoring system used to infer prognosis by assigning points for the following clinical factors: clinical signs, hepatic encephalopathy, ascites, hypoalbuminemia, hypoglobulinemia, hyperbilirubinemia, and prolongation of PTT) were calculated using patient data at the time of liver biopsy.^18^ Histopathology results, hepatic copper quantification via flame atomic absorption spectroscopy (ug/g dry weight liver)^a^, and aerobic/anaerobic culture results were recorded. Results of leptospirosis testing^b^ were documented, but testing was not required for inclusion.

The time point of initial presentation for liver biopsy was established as baseline (t=0), prior to the initiation of CsA. Values for serum ALP, ALT, AST, and GGT as well as serum albumin and total bilirubin were collected pre-treatment at t=0 and at sequential time points when the patient was rechecked. Since dogs were re-evaluated at variable time points, the data was categorized into seven time intervals after baseline 0: 1 (2-4 weeks), 2 (1-3 months), 3 (3-6 months), 4 (6-9 months), 5 (9-12 months), 6 (1-2 years), and 7 (> 2 years). If more than one dataset was available per time interval, the more complete and/or recent dataset was used. Due to variations in reference ranges between laboratories (Colorado State University Clinical Pathology Laboratory, Cummings Veterinary Medical Center at Tufts University Clinical Pathology Laboratory, IDEXX Laboratories, Antech Laboratories, or in-house referring veterinarian machines), each biochemical absolute value except for albumin, was converted to factor times the upper limit (xULN) of normal to standardize the data. Albumin values were converted to factor times the lower limit of normal (xLLN). The dose of CsA (mg/kg/day) and duration of therapy (months post initiation of cyclosporine) were documented at each time point. Clinical adverse effects suspected to be associated with CsA administration; such as inappetence, vomiting, diarrhea, gingival hyperplasia, secondary infections, hirsutism, papillomatosis, neoplasia, nephrotoxicity, hepatotoxicity, and diabetes mellitus were recorded. Hepatotoxicity or nephrotoxicity was defined as a 2-fold increase in ALT or creatinine compared to baseline, respectively, after the initiation of CsA therapy. If the liver or renal values improved/resolved without discontinuation of CsA, the case was not classified as hepatotoxicity or nephrotoxicity.

Remission of disease was defined as complete normalization of serum ALT to within the laboratory reference range (< 1.1 xULN).

Dogs that relapsed (ALT increased to > 1.1 xULN) upon tapering of CsA dosage were recorded and of those, dogs that achieved remission again after a dose increase were also recorded. Outcomes for each dog were categorized as alive, lost to follow up, death, or euthanized due to hepatic or non-hepatic disease. Outcome data for each dog was obtained via retrospective medical record review.

### Statistical analysis

Descriptive statistics were performed for the following variables: age (years), gender, breed, weight (kg), possible adverse effects of CsA, diet, concurrent medications, hepatic culture results, leptospirosis results, clinical score, presence of ascites, hypoalbuminemia, hyperbilirubinemia, hepatic copper concentrations (ug/g dry weight), PT/aPTT, pre-and post-treatment serum ALP, ALT, AST, GGT, total bilirubin (xULN), albumin (xLLN), dose of CsA at remission (mg/kg/day), duration of therapy till remission (months), attainment of remission, and sustained remission or relapse upon tapering CsA. Frequencies and percentages were calculated for categorical variables. Means and/or medians with standard deviations or ranges, respectively were calculated for continuous variables.

Continuous outcome liver parameter data for serum ALP, ALT, AST, GGT, bilirubin were assessed for normality with a Shapiro-Wilk test. If the data was not normally distributed, a log transformation was performed for statistical analysis. A linear regression model was used to compare the serum liver markers at each time interval to baseline (t= 0) and identify statistical significance (defined as p-value of < 0.05). Repeated measures on the same dog were taken into account while performing the analyses. The dose of CsA was added to the linear regression model to evaluate the separate effects of dose or time interval on each liver parameter. Type III p-values were reported to describe the aggregate effect of cyclosporine therapy across all time intervals for each liver parameter.

The impact of each clinical factor on the probability of remission was analyzed. A Fisher’s exact test was used to assess the impact of the categorical variables (hepatic copper >1000ug/g, gender, ascites, hypoalbuminemia, hyperbilirubinemia, prolongations in PT or PTT) on remission. A Wilcoxon Two-Sample test was used to examine the effect of non-parametric continuous variables; such as clinical scores and duration of therapy with remission. A Student’s t-test was used for assessment of parametric continuous variables (age, weight, and pre-treatment ALT [xULN]). Likelihood of remission was also analyzed with a Cox Proportional Hazards Model of survival analysis. Dogs that did not achieve remission or were lost to follow up were censored in the analysis. Hazard ratios were calculated for each clinical factor. A log-rank test was used to analyze the relationship between each categorical variable and time to remission. All statistical analyses were performed by a statistician using a commercial software package.^c^

## Results

### Patient population

Forty-eight dogs were included in the study of which 26/48 (54%) were female spayed, 20/48 (42%) were male castrated, and 2/48 (4%) were male intact. Median age was 8.5 years (range, 0.7-14 years). Median weight was 19.4 kg (range, 2.2-45kg). Twenty-two different breeds were represented: mixed breed (12), Labrador retriever (10), standard poodle (4), Brittany spaniel (2), Maltese (2), golden retriever (1), fox terrier (1), English springer spaniel (1), Scottish terrier (1), pug (1), miniature pinscher (1), Pomeranian (1), beagle (1), Havenese (1), Italian greyhound (1), beagle (1), Staffordshire terrier (1), basenji (1), wheaten terrier (1), bichon frise (1), ocherese (1), Yorkshire terrier (1), Doberman pinscher (1). Biopsies were performed laparoscopically (39/48), via ultrasound guidance (6/48), and surgically (3/48).

### Medications

All dogs were administered a microemulsified form of CsA^d^ approved for use in dogs. Only one dog was switched to an unknown brand generic CsA after the dog attained remission and the dog remained in remission even after the change in medication. Forty-three of 48 (90%) dogs were on concurrent hepatoprotectants; such as ursodiol (60%, 29/48), s-adenosylmethionine^e^ (79%, 38/48), vitamin E (29%, 14/48), and/or silybin (2%, 1/48). Nineteen of 48 dogs (40%) received a course of antibiotic therapy (combinations or monotherapy of amoxicillin, clindamycin, amoxicillin/clavulanic acid, metronidazole, and/or enrofloxacin) prior to or shortly after the liver biopsy was performed. Twelve dogs (25%) were on prednisolone prior to CsA therapy and were tapered off corticosteroids completely before (2/12) or within 6 months of CsA initiation (10/12). Of the 12 dogs, one was on azathioprine concurrently, which was tapered and discontinued before starting CsA. Another dog was on mycophenolate for immune-mediated polyarthritis when she was diagnosed with CH, but the drug was ultimately tapered and discontinued. One dog received a single dose of dexamethasone in the hospital. Seven dogs were treated with penicillamine prior to starting CsA. One dog was started on losartan and one on colchicine. Three dogs were on therapy for ascites (spironolactone +/− furosemide).

### Concurrent diet

Twenty of 46 dogs (43%) with available diet information were on commercially available prescription hepatic diets^f^ (19/20) or home-cooked low copper diets (1/20) at the time of diagnosis.

### Liver culture results

Four of 40 (10%) of aerobic liver cultures were positive. Organisms grown were Acinetobacter (1/4), Streptococcus beta hemolytic (1/4), Staphylococcus (1/4), and Staphylococcus coagulase negative (1/4). One of the 37 (2.7%) anaerobic liver cultures was positive (Clostridium septicum). Only one dog had a bile culture performed and results were negative for aerobic and anaerobic growth.

### Leptospirosis results

Seventeen of 48 dogs (35%) had leptospirosis microscopic agglutination serology performed of which 4 showed seroreactivity. Three of the 4 dogs had recent vaccinations and all of these dogs failed to have improvement in serum liver enzyme values with treatment with ampicillin or doxycycline. Likewise, the fourth dog with positive seroreactivity (but with no history of vaccination) had no response to a prolonged course of doxycycline.

### Liver parameter changes and remission with CsA treatment

Thirty eight of 48 dogs (79%) attained biochemical remission with a normalization of serum ALT. Median starting dose of CsA was 7.8 mg/kg/day (range, 1.5-12.7mg/kg/day) and median dose at the time of remission was 7.9 mg/kg/day (range, 2.5-12.7mg/kg/day). The median time to remission was 2.5 months (range, 0.75-12 months). After remission was reached, 22 of 33 (67%) dogs experienced a relapse when the CsA dose was tapered. The tapering protocol was not standardized from dog to dog. Nine of these 22 dogs achieved remission again when the CsA dose was increased. One dog was tapered off cyclosporine completely and remained in remission 12 months after discontinuation of the drug.

Median absolute values and ranges for ALP, ALT, AST, GGT, bilirubin, and albumin in response to CsA treatment in the 7 defined time ranges are shown in Table 1 and all the liver parameters except albumin showed significant decreases with cyclosporine therapy. The fold increases over the ULN for serum ALT and AST were significantly decreased at post treatment intervals 1-7 (P< 0.003) in comparison to baseline (Figure 1). The fold increases over the ULN for ALP and GGT were significantly decreased at post treatment intervals 2-7 (P < 0.0001) and interval 3-7 (P < 0.0001) respectively (Figure 2). The fold increases in bilirubin were significantly decreased for time intervals 2-7 (P<0.004). P values were not significant for xLLN albumin at any time interval (Figure 3).

**Table 1:** Changes in each biochemical liver parameter at each time interval post CsA therapy. Medians and ranges for biochemical parameters at each of the seven time intervals post initiation of cyclosporine (CsA) treatment. The * indicates statistical significance (p-value of < 0.05) when comparing the fold increase over the ULN (for ALT, AST, GGT, ALP, TB) or the fold decrease from the LLN (for albumin) at each post-treatment time interval to baseline (t = 0). A type III p-value < 0.05, labeled with superscript a, indicates an overall statistically significant decrease in liver values in response to CsA. After controlling for CsA dose, an adjusted type III p-value < 0.05, labeled with superscript b, denotes an overall significant decrease in liver values in response to CsA if CsA dose was held constant. N is the number of dogs with available liver parameter data at each time interval.

|  | Time Interval |  |  |  |  |  |  |  |  |  |
| --- | --- | --- | --- | --- | --- | --- | --- | --- | --- | --- |
|  | 0 | 1 | 2 | 3 | 4 | 5 | 6 | 7 |  |  |
|  | 0 | 2-4 weeks | 1-3 months | 3-6 months | 6-9 months | 9-12 months | 1-2 years | > 2 years | Type 3 p-value | Controlling for dose type 3 p-value |
| ALT (IU/L) | 615<br>(185-3015)<br>n = 48 | 273<br>(32-4984)*<br>n = 32 | 114<br>(15-1645)*<br>n = 36 | 105<br>(27-960)*<br>n = 35 | 86<br>(22-312)*<br>n = 31 | 86<br>(21-464)*<br>n = 25 | 81<br>(21-1171)*<br>n = 27 | 63<br>(19-443)*<br>n = 19 | <0.0001<sup>a</sup> | <0.0001<sup>b</sup> |
| AST (IU/L) | 115<br>(37-480)<br>n = 42 | 50<br>(17-1491)*<br>n = 22 | 33<br>(17-246)*<br>n = 26 | 36<br>(11-186)*<br>n = 27 | 29<br>(18-181)*<br>n = 24 | 33<br>(24-96)*<br>n = 17 | 35<br>(15-123)*<br>n = 23 | 29<br>(17-103)*<br>n = 14 | <0.0001<sup>a</sup> | 0.003<sup>b</sup> |
| ALP (IU/L) | 375<br>(61-5236)<br>n = 48 | 247<br>(30-5357)*<br>n = 30 | 130<br>(20-1746)*<br>n = 35 | 125<br>(24-2205)*<br>n = 33 | 89<br>(15-431)*<br>n = 29 | 81<br>(22-638)*<br>n = 23 | 110<br>(22-5821)*<br>n = 27 | 114<br>(29-638)*<br>n = 19 | <0.0001<sup>a</sup> | 0.004<sup>b</sup> |
| GGT (IU/L) | 16<br>(0-316)<br>n = 42 | 12<br>(0-302)<br>n = 23 | 5.5<br>(0-87)*<br>n = 24 | 5<br>(0-50)*<br>n = 25 | 3<br>(0-43)*<br>n = 23 | 5<br>(0-16)*<br>n = 19 | 2.5<br>(0-229)*<br>n = 22 | 2.5<br>(0-21)*<br>n = 16 | 0.002<sup>a</sup> | 0.004<sup>b</sup> |
| Total bilirubin (mg/dL) | 0.3<br>(0.1-14.7)<br>n = 46 | 0.5<br>(0-2.6)<br>n = 26 | 0.3<br>(0.1-0.8)*<br>n = 29 | 0.2<br>(0.1-0.9)*<br>n = 31 | 0.2<br>(0.1-1)*<br>n = 25 | 0.2<br>(0.1-0.4)*<br>n = 20 | 0.1<br>(0-1.8)*<br>n = 27 | 0.1<br>(0.1-1.5)*<br>n = 18 | 0.004<sup>a</sup> | 0.003<sup>b</sup> |
| Albumin (g/dL) | 3.3<br>(1.6-4.7)<br>n = 44 | 3.1<br>(1.8-3.9)<br>n = 27 | 3.2<br>(2.5-4.3)<br>n = 32 | 3.3<br>(2.2-4)<br>n = 31 | 3.3<br>(2-4.3)<br>n = 26 | 3.1<br>(2-4.1)<br>n = 21 | 3.2<br>(2.2-3.9)<br>n = 27 | 3.2<br>(2.4-3.7)<br>n = 18 | 0.21 | 0.09 |

**Figure 1:**
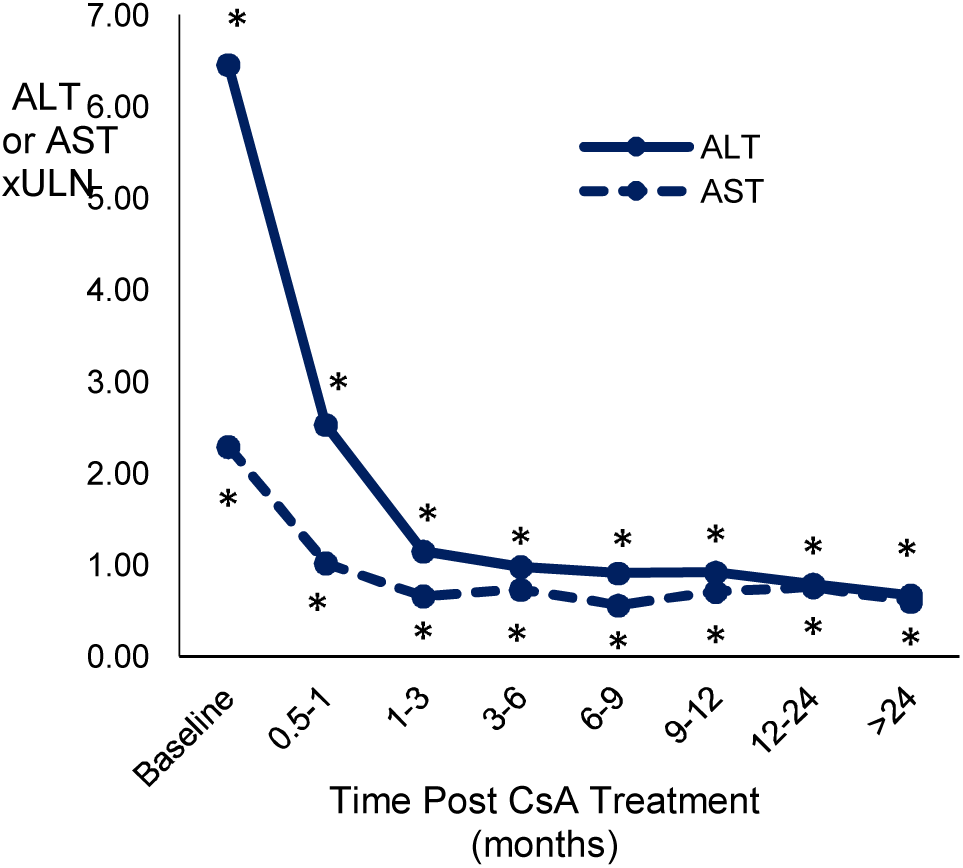
Change in serum ALT and AST activity in response to CsA therapy over time. Median increase over the upper limit of the reference value (xULN) for serum ALT and AST activity at different time intervals after starting CsA (cyclosporine) therapy. * = the median value of ALT or AST at that time interval was significantly reduced compared to the baseline value. ALT is depicted in the solid line and AST in the dashed line.

**Figure 2:**
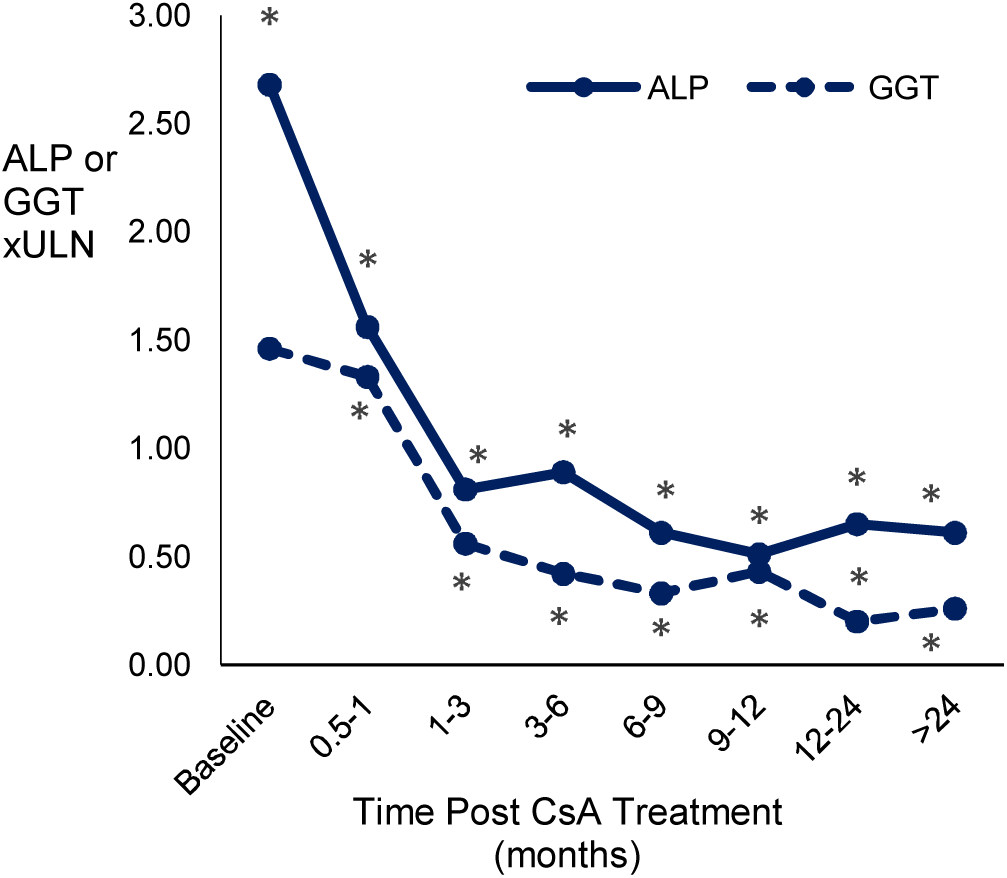
Change in serum ALP and GGT activity in response to CsA therapy over time. Median increase over the upper limit of the normal reference range (xULN) for serum ALP and GGT activity at different time intervals after starting CsA (cyclosporine) * = the median value of ALP or GGT activity at that time interval was significantly decreased compared to the baseline value. ALP is depicted in the solid line and GGT in the dashed line.

**Figure 3:**
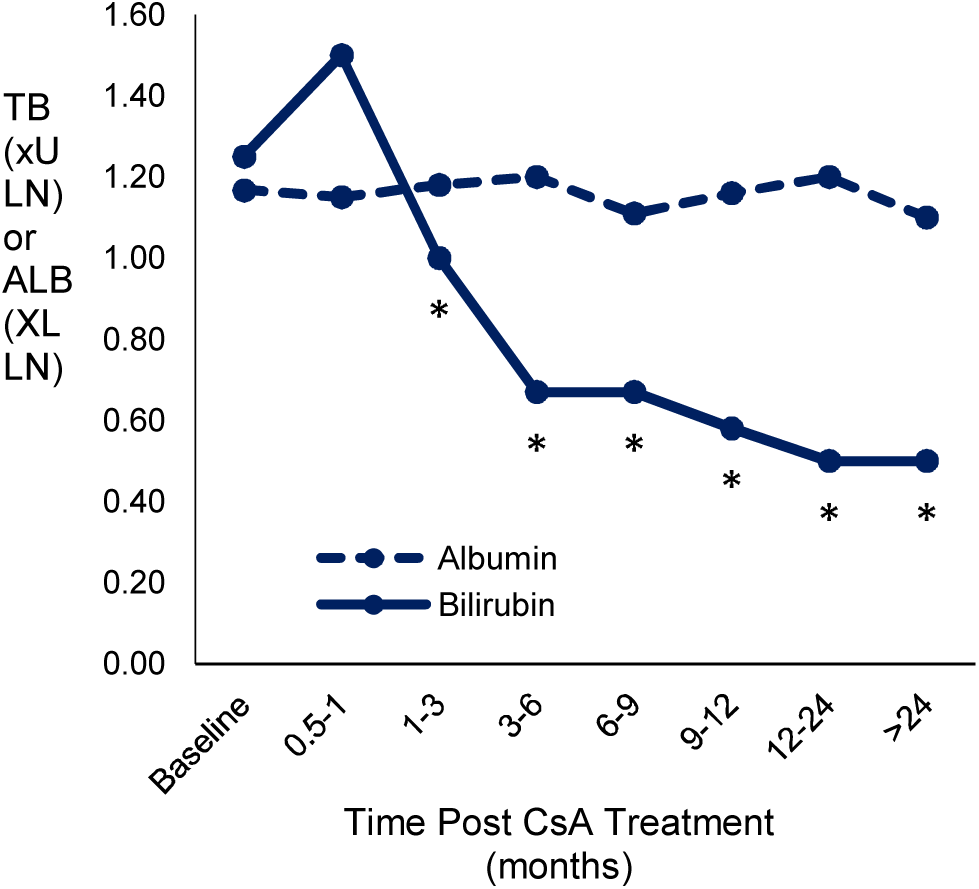
Changes in serum albumin and bilirubin in response to CsA therapy over time. Median decrease from the lower limit of the reference range (xLLN) for serum albumin (ALB) and the median increase over the upper limit of the reference range (xULN) for serum total bilirubin (TB) at post-treatment time intervals after starting CsA (cyclosporine). * = the median values for ALB were significantly decreased compared to baseline values. TB is depicted in the solid line and ALB in the dashed line.

Type III P-values showed the overall effect of CsA on each liver parameter. The fold increases over the ULN for serum ALP, ALT, and AST (type III P < 0.0001) and for serum GGT (type III P = 0.002) were significantly decreased post-treatment compared to baseline, but were not significantly different for xLLN albumin (type III P = 0.21). When CsA dose was held constant to remove the effect of this variable, the fold increases over ULN for serum ALP, ALT, AST, and GGT remained significantly decreased post-treatment (Type III P-values in Table 1).

### Clinical factors and their effect on attaining remission

No clinical or biochemical factor examined was significantly associated with remission (Table 2). Fisher’s exact tests and Cox Proportional Hazards model of survival analysis demonstrated that the probability of attaining remission was not impacted by ascites, hyperbilirubinemia, hepatic Cu > 1000ug/g, or prolongation of PT/aPTT (Table 2). Median clinical score was 1.5 (range 0-7). Clinical score did not significantly differ between dogs that achieved remission and those that did not (p = 0.61). Institution (p = 0.71), age (p = 0.87), gender (p = 0.83), weight (p = 0.48), pre-treatment xULN ALT (p = 0.67), duration of therapy (p = 0.74), and starting CsA dose (p = 0.50) did not significantly impact the likelihood of remission.

**Table 2:** Association of prognostic clinical factors with the likelihood of attaining remission with CsA. The p-values indicate the association of each clinical factor with likelihood of attaining remission. Fisher’s exact tests and Cox proportional hazards model of survival analyses were performed. No factor examined was significantly associated with remission. The number and percentage of dogs with each clinical factor is listed. The number of dogs with each clinical factor that attained remission is also shown.

| **Prognostic Clinical Factors** | **% of dogs with Clinical Factor** | **% which attained remission** | **Fisher's exact test p value** | **Survival Analysis p-value** |
| --- | --- | --- | --- | --- |
| Ascites | 8/47 (17%) | 6/8 (75%) | 1.00 | 0.27 |
| Hypoalbuminemia | 6/47 (13%) | 5/6 (83%) | 1.00 | 0.11 |
| Hyperbilirubinemia | 25/47 (53%) | 20/25 (80%) | 1.00 | 0.54 |
| Prolonged PT or aPTT | 6/41 (15%) | 5/6 (83%) | 1.00 | 0.71 |
| Hepatic Cu > 1000ug/g | 11/45 (24%) | 8/11 (73%) | 0.69 | 0.86 |

### Clinical factors and their effect on time to remission

Log-rank analysis showed that neither starting dose of CsA nor the following clinical factors impacted time to remission (ascites p-value = 0.22, hypoalbuminemia p-value = 0.08, hyperbilirubinemia p-value = 0.51, hepatic Cu > 1000ug/g p-value = 0.85, prolonged PT/aPTT p-value = 0.68).

### Adverse effects

The most common adverse effects associated with CsA therapy were gastrointestinal signs in 18/48 (38%) including hyporexia, vomiting, and/or diarrhea (Table 3). Seventeen of 18 dogs experienced gastrointestinal signs that were transient and improved with a decrease in dose or frequency of administration (8/17), freezing the capsule (3/17), adaptation to the CsA treatment over time (within 2 months) (5/17), administering with food (1/17). The second most common side effect was gingival hyperplasia, which occurred in 12/48 (25%) of dogs. Less common potential side effects included opportunistic infections (4/48), lymphoma (3/48), acute kidney injury (3/48), hirsutism (2/48), papillomatosis (1/48), diabetes mellitus (1/48), and head tremors (1/48). The opportunistic infections observed were urinary tract infections (2: *E.coli* and *Proteus* infection), pyoderma (1) with a Bacillus sp. (grown on enrichment broth only), and ringworm (specific species unknown) (1).

**Table 3:**
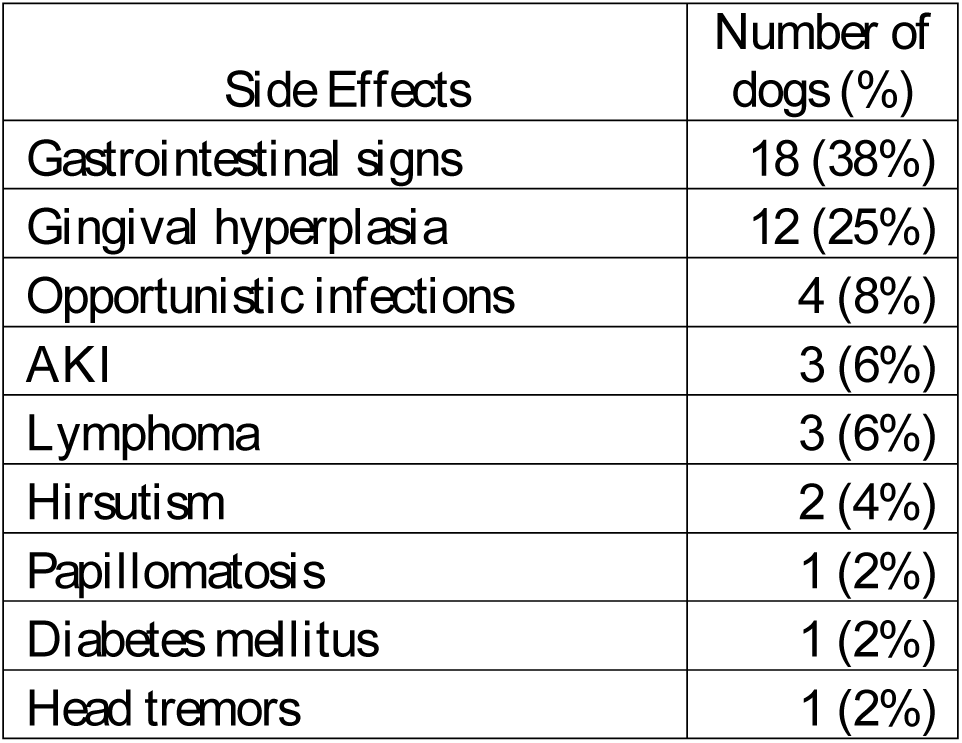
Potential adverse effects associated with CsA administration

Three dogs were diagnosed with lymphoma. One had leukemic lymphoma with metastasis to the liver, spleen, heart, lymph nodes, lung, and glomeruli on necropsy 8 months after starting CsA. The second dog had lymphoblasts on analysis of peritoneal effusion 19 months after starting CsA. The third dog was diagnosed with multicentric lymphoma 6 months after starting CsA. Cyclosporine doses at the time of lymphoma diagnosis were 4.5mg/kg/day, 2.9mg/kg/day, and 3.7mg/kg/day, respectively for each dog. All dogs had evidence of lymphoplasmacytic infiltrates on their initial liver biopsies, but no evidence of lymphoma.

Two dogs experienced acute kidney injury 2 and 4 years after being on CsA, respectively. In both dogs, an etiology was not identified, but in the first dog, a necropsy showed severe chronic interstitial nephritis and fibrosis with glomerulosclerosis.

Cyclosporine was discontinued in 3 dogs for vomiting [1], suspect bronchopneumonia [1], and foaming at the mouth when receiving the liquid cyclosporine [1].

### Outcome

Survival outcome at the last available time-interval (range, 24-65 months) were as follows: alive and doing well (21/48), lost to follow-up (10/48), euthanized due to non-hepatic disease (13/48), euthanized due to hepatic disease (2/48), or died/euthanized for an unknown cause (2/48). Three dogs were necropsied after euthanasia due to non-hepatic disease. Two dogs had evidence of hepatic fibrosis (mild to moderate in one, marked in the other) and nodular regeneration, but there was no inflammatory infiltrate on liver histopathology after 16 months and approximately 4 years of CsA therapy. The third dog had mild multifocal accumulations of lymphocytes and neutrophils. None of the dogs that were necropsied had normal ALT at the time of euthanasia.

## Discussion

In the current retrospective case series, remission, defined as normalization of serum ALT, was attained in 79% of dogs treated with CsA for CH. The median duration of therapy necessary to obtain remission was 2.5 months (0.75-12 months) and the median dose in the dogs that attained remission was 7.9 mg/kg/day (2.5-12.7mg/kg/day). Significant improvements were seen in response to CsA for serum ALP, ALT, AST, GGT, and total bilirubin. None of the clinical features and biochemical parameters previously associated with a poor prognosis in CH, such as; ascites^6,41^, hypoalbuminemia, ^6,21^ hyperbilirubinemia,^6,42^ prolonged PT/aPTT, ^6,18,21^ or higher clinical score^18^ negatively impacted the likelihood of remission or time to remission. Side effects of CsA were minimal. The most common side effects were gastrointestinal upset and gingival hyperplasia. The results of this study provide preliminary evidence that CsA is a well-tolerated and effective therapy for dogs with suspected immune mediated CH.

None of the previously identified poor prognostic factors associated with CH^6, 18, 21, 41,42^ were found to impact the ability of CsA to induce remission, indicating that treatment of CH with CsA may be successful even in dogs with later stage disease. Although serum albumin, which is considered a marker of more advanced disease, did not change over time in this study, this may have been a type II error due to small sample size and the mild degree of hypoalbuminemia. Hypoalbuminemia was present in only 6 dogs in this study of which only 2 had markedly decreased albumin levels (1.6 g/dL and 1.8 g/dL). Although not statistically significant, serum albumin levels did normalize with CsA therapy in 5 of the 6 hypoalbuminemic dogs, including the two with markedly decreased values.

The failure of remission in 21% of the dogs in this study could be explained by the individual variability in the pharmacology and pharmacodynamic response to CsA that is known to exist in dogs.^24^ Intrinsic differences of CsA among dogs are associated with several factors including variations in cytochrome p450 activity, MDR1 status, the extent of protein binding, and the drug’s volume of distribution.^24^ Other factors that could have impacted the pharmacokinetics of CsA in this study include administration of CsA with food,^43,44^ concurrent medications,^45–47^, freezing the medication,^48^ and the CsA dose. A previous study has demonstrated that administration of CsA with food decreases the drug’s bioavailability,^43^ while a separate study found administration of CsA with food had no effect on the efficacy of the drug in dogs.^44^

Freezing the CsA capsule for 28 days has not been shown to affect integrity of the capsule, absorption of the medication, or plasma concentrations;^48^ however, the dogs in that study were healthy research animals in contrast to animals with hepatic disease.^49^ Performing pharmacologic or pharmacodynamic testing; such serum CsA levels or T-cell inhibition assays^50,51^ would help elucidate the impact of the such factors on the bioavailability and immunosuppressive effects of CsA. Future studies examining the relationship of CsA dosing to serum levels and pharmacodynamic response (T-cell inhibition) are needed in dogs with hepatic disease. These studies might help to define the minimally effective dose and improve response to therapy in dogs with CH.

Other potential explanations for failure to attain remission with CsA therapy are owner noncompliance, the presence of comorbidities such as gastrointestinal/endocrine disease, lack of therapeutic efficacy, or concurrent medications that induce serum liver enzyme concentrations. Alternatively, dogs that failed to respond to CsA may not have had immune mediated CH.

In human medicine, autoimmune hepatitis is a well described condition with established diagnostic criteria and recommendations for therapy.^40^ Response to immunosuppressive therapy is included in the diagnostic algorithm for immune hepatitis in people.^40^ Prednisone and azathioprine are first line therapies, while CsA and mycophenolate are used as rescue agents or in cases refractory to prednisone and azathioprine.^39,40^ There is no standardized criteria for the diagnosis of autoimmune hepatitis in dogs or consensus on the best therapeutic approach for suspected immune hepatitis in dogs. Two retrospective studies looking at the use of corticosteroids to treat CH in dogs have suggested that some dogs do have a positive response to therapy and thus meet one of the diagnostic criteria for immune mediated disease.^21,22^ The response in this study to CsA, a T-cell inhibitor with more targeted immunosuppressive activity, further bolsters this hypothesis.

Although previous studies suggested that corticosteroids are beneficial in some dogs with CH,^21,22^ there are a number of associated side effects with corticosteroids,^23^ including the induction of serum liver enzyme activity (ALP > GGT >>> ALT) and the development of a vacuolar hepatopathy.^52^ In one study of corticosteroid use in dogs with CH, 15/36 dogs treated with prednisolone had a vacuolar hepatopathy on re-biopsy.^21^ Although typically a benign morphologic change, vacuolar hepatopathies can rarely be associated with functional hepatic disease.^52^ Corticosteroids also have the potential to worsen hepatic encephalopathy,^53^ facilitate sodium retention to exacerbate ascites,^54^ and promote hypercoagulability predisposing to thrombosis.^55^ In some cases, corticosteroids have shown to be of no benefit and indeed may be harmful in dogs with late-stage disease and cirrhosis.^22,56^ These hepatic consequences of corticosteroid use can be averted with the administration of CsA. Additionally, the use of CsA makes it easier to use serum liver enzyme activity as a biomarker of response to therapy.

This study also showed that CsA was fairly well tolerated. The most common side effects were gastrointestinal signs, which were typically transient. Multiple publications have documented that gastrointestinal signs are the most common side effects in dogs treated with CsA for atopic dermatitis.^25–28^ Gingival hyperplasia was the second most common side effect observed in our study, but typically can be managed with azithromycin or gingivectomy if needed.^57,58^ Urinary tract infections and opportunistic infections have also been infrequently observed with CsA therapy.^33–37^ More severe adverse events seen in this retrospective; such as AKI or lymphoma have an uncertain association with CsA use. These events occurred at least 2 years after the initiation of CsA making it difficult to establish causality. There is only one report of nephrotoxicity associated with CsA overdose in a clinical^59^ setting and one report of multicentric lymphoma associated with CsA administration.^60^ No cases of hepatotoxicity were identified in this study although experimentally, CsA can cause liver injury.^61^

There were 4 cases with positive liver cultures. These cases were treated with appropriate antibiotics; however, remission was not achieved with antibiotic therapy alone. These cases eventually responded to CsA administration, which suggests that the primary etiology was immune-mediated and not infectious. In some cases, these bacteria could have been contaminants or opportunistic infections rather than primary pathogens. In future studies, fluorescent in situ hybridization staining of the liver could elucidate active involvement of bacterial pathogens in dogs with hepatitis if liver/bile cultures are positive.

## Limitations

This study had several limitations.

Although established WSAVA criteria^8^ were used for histopathologic diagnosis of CH and all biopsy reports were reviewed by a board-certified internist with expertise in hepatology, all the liver biopsies were not interpreted by the same boarded histopathologist, which may have introduced variability in the final diagnosis. Due to this limitation, we did not assess the effect of stage (severity of fibrosis) and grade (inflammation and degenerative change) on histopathology with treatment response. Variable reference ranges existed amongst laboratories at the two academic institutions and send-out or in-house biochemistry machines. Therefore, liver values were converted to xULN or LLN in an attempt to standardize the data.

In humans, the diagnostic criteria for autoimmune hepatitis are clearly established and scored based on the following features: elevated serum ALT, the presence of positive serum autoantibodies titers, elevated serum immunoglobulin G, characteristic histopathology findings (interface hepatitis), and the exclusion of other etiologies such as viral and alcoholic hepatitis.^39,40^ Although medical records were comprehensively and expertly reviewed, it is possible given the retrospective nature of the study that some cases were not truly idiopathic CH. Remission criteria are also standardized in humans and include normalization of serum ALT, bilirubin and IgG, alleviation of clinical symptoms, and return to normal hepatic tissue on histopathology.^40^ Criteria for the diagnosis of immune hepatitis in dogs and what constitutes remission have not been established for dogs.^62^ We chose to use normalization of serum ALT to define remission; however, studies have shown that serum ALT in the dog does not have the sensitivity to detect subtle changes in CH.^7^ Ideally, in addition to biochemical remission, we would have also evaluated clinical remission (resolution of clinical signs) and histological resolution of CH (based on examination of repeat liver biopsies).

Since this was a retrospective study, treatment and monitoring protocols were not standardized. Dogs were administered CsA at variable dosing and tapering schedules. Furthermore, the majority of the dogs in the study received concurrent hepatoprotective medications, which may have improved liver enzyme concentrations.^63–65^

Recheck appointments were scheduled at different time points and clinics at the discretion of the primary clinician and/or client. As a result, long-term follow-up was limited. Eleven of 48 dogs had unknown outcome data and only nineteen dogs had available serum ALP and ALT data > 2 years after baseline. Due to this lack of follow-up, survival analysis could not be performed. Additionally, there may have been selection bias towards dogs with more positive outcomes. The exclusion of dogs on CsA for less than 2 weeks and the selection of dogs at tertiary academic institution referral hospitals may have contributed to this bias.

## Conclusion

In conclusion, this study demonstrated that CsA is a well-tolerated and potentially effective medication in normalizing serum liver enzymes and total bilirubin in dogs with idiopathic CH. Previously reported poor prognostic factors in CH did not affect the ability of CsA to induce remission. These findings provide further evidence that in some dogs, CH is immune-mediated. Further prospective studies will be necessary to determine whether CsA therapy results in histopathologic remission of disease and to compare CsA with other immunomodulatory medications to determine which therapy is safer and more effective in attaining remission and extending survival in dogs with CH.

## Footnote

a Colorado State University Diagnostic Laboratory, Fort Collins, USA

b Colorado State University Diagnostic Laboratory or IDEXX

c SAS v9.4 (SAS Institute Inc., Cary, NC)

d Nutramax Laboratories

e Atopica^®^, Elanco Animal Health

f Hill’s l/d or Royal Canin Hepatic

## Acknowledgements

Conflict of Interest Declaration: Authors declare no conflict of interest.

Off-label Antimicrobial Declaration: Authors declare no off-label use of antimicrobials.

## Abbreviations

CH: chronic hepatitis
CsA: cyclosporine
WSAVA: World Small Animal Veterinary Association
ALP: alkaline phosphatase
ALT: alanine aminotransferase
AST: aspartate aminotransferase
GGT: gamma-glutamyl transferase
PT: prothrombin time
aPTT: activated partial thromboplastin time
xULN: factor times upper limit of normal
xLLN: factor times lower limit of normal

